# Modifying TIMER, a slow-folding DsRed derivative, for optimal use in quickly-dividing bacteria

**DOI:** 10.1101/2021.01.12.426338

**Authors:** Pavan Patel, Brendan J. O’Hara, Emily Aunins, Kimberly M. Davis

**Affiliations:** W. Harry Feinstone Department of Molecular Microbiology and Immunology, Johns Hopkins Bloomberg School of Public Health, Baltimore, MD, USA; Department of Molecular Biology and Microbiology, Tufts University School of Medicine, Boston, MA, USA

## Abstract

It is now well appreciated that members of pathogenic bacterial populations exhibit heterogeneity in growth rates and metabolic activity, and it is known this can impact the ability to eliminate all members of the bacterial population during antibiotic treatment. It remains unclear which pathways promote slowed bacterial growth within host tissues, primarily because it has been difficult to identify and isolate slow growing bacteria from host tissues for downstream analyses. To overcome this limitation, we have developed a novel variant of TIMER, a slow-folding fluorescent protein, to identify subsets of slowly dividing bacteria within host tissues. The original TIMER folds too slowly for fluorescence accumulation in quickly replicating bacterial species (*Escherichia coli, Yersinia pseudotuberculosis*), however this TIMER_42_ variant accumulates signal in late stationary phase cultures of *E. coli* and *Y. pseudotuberculosis*. We show TIMER_42_ signal also accumulates during exposure to sources of nitric oxide (NO), suggesting TIMER_42_ signal detects growth-arrested bacterial cells. In a mouse model of *Y. pseudotuberculosis* deep tissue infection, TIMER_42_ signal is clearly detected, and primarily accumulates in bacteria expressing markers of stationary phase growth. There was not significant overlap between TIMER_42_ signal and NO-exposed subpopulations of bacteria within host tissues, suggesting NO stress was transient, allowing bacteria to recover from this stress and resume replication. This novel TIMER_42_ variant represents a new faster folding TIMER that will enable additional studies of slow-growing subpopulations of bacteria, specifically within bacterial species that quickly divide.

**Author Summary:** We have generated a variant of TIMER that can be used to mark slow-growing subsets of *Yersinia pseudotuberculosis*, which has a relatively short division time, similar to *E. coli*. We used a combination of site-directed and random mutagenesis to generate the TIMER_42_ variant, which has red fluorescent signal accumulation in post-exponential or stationary phase cells. We found that nitric oxide (NO) stress is sufficient to promote TIMER_42_ signal accumulation in culture, however within host tissues, TIMER_42_ signal correlates with a stationary phase reporter (*dps*). These results suggest NO may cause an immediate arrest in bacterial cell division, but during growth in host tissues exposure to NO is transient, allowing bacteria to recover from this stress and resume cell division. Thus instead of indicating a response to host stressors, TIMER_42_ signal accumulation within host tissues appears to identify slow-growing cells that are experiencing nutrient limitation.

## Introduction

Identifying differences in the growth rates of individual cells can provide significant information in many biological systems. Quickened growth rates can indicate transformative properties in mammalian cells, indicate that specific B cell populations are expanding in response to infection or vaccination, or suggest distinct differentiation programs are occurring within cell populations during the developmental stages of an organism. For populations of bacterial cells, rapidly dividing bacteria are generally more metabolically active and thereby more susceptible to antibiotic treatments, whereas slowly dividing cells have decreased metabolic activity and are less susceptible to antibiotic treatment (1-3).

Many research groups have begun to develop tools to differentiate between faster and slower dividing cells, which enable downstream studies to identify additional differences within subpopulations. Many of these tools, such as slow-folding fluorescent proteins, were initially developed to study eukaryotic systems (4, 5). However, several of these approaches have been adapted to study bacterial populations, primarily to better understand the pathways that lead to the formation of slow-growing subsets of bacteria with reduced antibiotic susceptibility. Some bacterial tools to detect differential cell division rates include: fluorescence dilution or plasmid loss approaches (6-8); detection of DNA replication machinery (9, 10); and differential labeling of the cell envelope to detect newly synthesized macromolecules (11, 12); which can all be used in combination with fluorescence microscopy and live imaging techniques. It has been difficult to apply these techniques to identify slow growing bacteria in the context of mammalian infection models, since many infection models are not amenable to live imaging platforms, and instead utilize endpoint analyses. Bacterial population numbers can also contract and expand dramatically during infection (13, 14), which can dilute fluorescent signals or plasmids very early after inoculation and result in a complete loss of signal.

Slow-folding fluorescent proteins may offer an alternative method to identify slow growing cells within bacterial populations. TIMER is a slow-folding derivative of the tetrameric fluorescent protein, DsRed, that has been successfully used to detect differential replication rates of *Salmonella enterica* serovar Typhimurium (*S*. Typhimurium) during growth in the spleen (15). Wild-type (WT) DsRed generates both green and red monomers during fluorescence maturation; green monomers fold more quickly than red monomers, which require an extra amidation reaction during folding (16). The green monomeric species is also less stable, resulting in incorporation of very few (∼10%) green monomers into tetrameric DsRed, and predominantly red signal (16). In contrast, TIMER contains a S197T mutation that stabilizes the green monomeric form (4, 17). When TIMER is initially expressed, this results in an early accumulation of predominantly green monomers and green fluorescent signal. With continued translation and folding, red monomeric species are incorporated into newly formed tetramers, which generates mixed ‘orange’ tetramers containing both green and red monomers (17). In fast-growing cells, cell division occurs before red monomers fold, and green signal predominates. In slow-growing cells, red monomers are incorporated before cell division occurs; therefore mixed green and red tetramers, can be detected. The accumulation of red fluorescent signal requires TIMER protein folding prior to cell division (15), and so the utility of this tool is dependent on the cell division rate of the cells of interest. Red TIMER signal emerges within *S*. Typhimurium dividing approximately 0.16 times/hour, or one doubling every 6 hours (15), but as we show here, this original TIMER variant folds too slowly for signal detection in more quickly dividing bacteria, such as *Escherichia coli* or *Yersinia pseudotuberculosis*. Each of these species can divide as quickly as 1.5-2 times/hour, or one doubling every 30-45 minutes, in culture (18, 19).

*Yersinia pseudotuberculosis* in an enteric human pathogen that can escape the intestinal tract and replicate within deep tissue sites to form clonal extracellular clusters of bacteria called microcolonies (or pyogranulomas) (20-22). Multiple immune cell types are recruited to the site of infection; however, bacteria continue to replicate, aided by their large arsenal of virulence factors (23-25). Bacteria at the periphery of microcolonies respond to neutrophil contact by expressing high levels of the anti-phagocytic type-III secretion system and its associated effector proteins (22). Peripheral bacteria also respond to and detoxify the diffusible antimicrobial gas, nitric oxide (NO), which is produced by recruited monocytes circumscribing the neutrophil layer (22, 26). NO detoxification occurs through expression of *hmp*, a nitric oxide dioxygenase, which converts antimicrobial NO to the innocuous NO_3_ (27). It remains unclear if the protective expression of *hmp* by peripheral cells comes at a fitness cost, and if this stress promotes the formation of a slow-growing subset of bacteria at the periphery of microcolonies. The preferential survival of *hmp-*expressing cells during antibiotic treatment supported this hypothesis (28), however our results here will suggest that NO exposure occurs transiently within host tissues, and that Hmp^+^ cells retain the ability to divide.

Here, we sought to generate a new TIMER variant to detect differential growth rates of *Y. pseudotuberculosis* (Yptb) within host tissues using both site-directed and random mutagenesis. This TIMER variant, TIMER_42_, accumulates red fluorescent signal late in stationary phase Yptb, allowing us to detect slow-growing subsets of bacterial cells during growth in culture and within host tissues. TIMER_42_ lacks green fluorescent signal, which allows us to compare red fluorescence accumulation to green fluorescent stress reporter constructs and ask if stressed cells have a heightened TIMER_42_ expression. Using these constructs together, we show that NO stress is sufficient to promote TIMER_42_ signal accumulation in culture; however, NO exposure within host tissues appears to be transient, and does not cause prolonged slowed growth. Instead, we detected a separate population of TIMER_42_^+^ bacteria within host tissues, which also have markers of stationary phase cells (*dps* expression). TIMER_42_ will be a very useful tool for future studies by enabling isolation of slow growing subsets of bacteria from host tissues for downstream analyses.

## Results

### Site-directed and random mutagenesis were used to generate a novel TIMER variant, TIMER_42_

To test the hypothesis that responses to reactive nitrogen species (RNS) were sufficient to slow the growth of stressed cells, we sought to develop tools to specifically detect slow-growing subpopulations of bacteria within *Y. pseudotuberculosis* microcolonies. We focused our efforts on TIMER, since this fluorescent protein had been successfully used to identify and isolate slow-growing populations of *S. Typhimurium* from the mouse spleen (3, 5).

TIMER was inserted into the high copy vector, pUC18, downstream of the IPTG-inducible *P*_*tac*_ promoter, and transformed into *E. coli. E. coli* cultures were grown 24 hours (h) in the presence of IPTG to induce TIMER expression, and fluorescence was detected by plate reader using 480nm excitation and emission detection from 500nm to 600nm to detect the incorporation of both green and red monomers into TIMER tetramers (15). Fluorescence was not detected in exponential phase cells (4h growth), and faint green signal could be detected from bacteria late in stationary phase (24h growth). However, this green signal remained below the background signal from non-fluorescent *E. coli*, and red TIMER signal did not accumulate **(Fig 1A)**. We then generated a TIMER_I161T T197A_ variant, which should lack green fluorescent signal and accumulate red signal more quickly due to quickened folding kinetics (1). We were hoping faster folding would allow us to detect red TIMER_I161T T197A_ fluorescence accumulation in post-exponential or stationary phase cells, and eliminating the green monomeric species would allow us to use green fluorescent stress reporters alongside TIMER_I161T T197A_. However, TIMER_I161T T197A_ also did not have detectable red fluorescent signal after 24 h growth with IPTG induction **(Fig 1A)**. Additionally, we quantified expression of TIMER_I161T T197A_ in a low copy vector, pMMB67EH, which we typically use for reporters in our mouse model of infection. We did not observe detectable fluorescence in stationary phase (24h growth) *E. coli* or *Y. pseudotuberculosis* (Yptb), in the presence or absence of IPTG induction **(Supplemental Fig 1)**.

**Fig 1:**
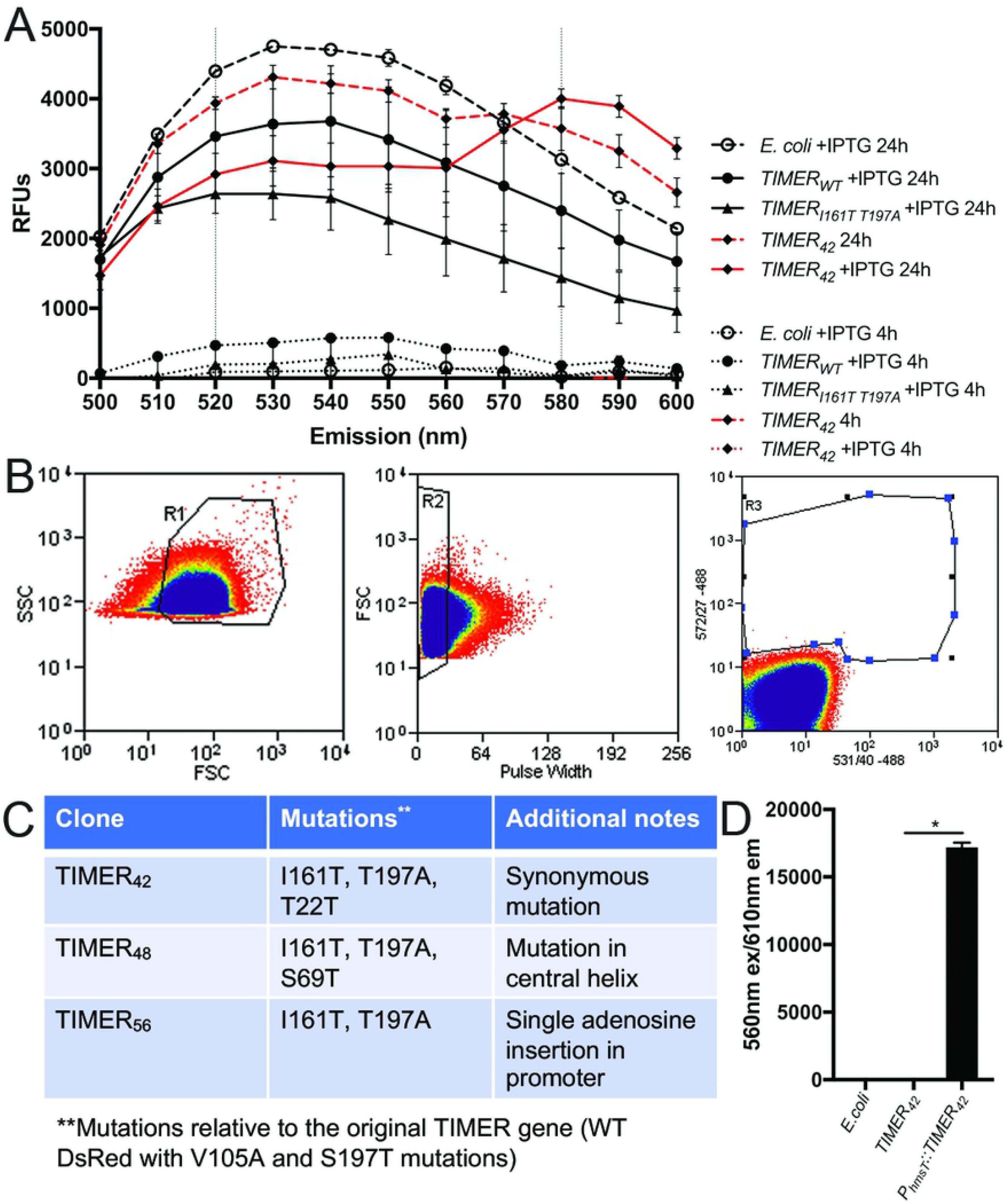
Site-directed and random mutagenesis were used to generate a novel TIMER variant, TIMER_42_. A) Clones were grown in LB in the presence and absence of IPTG for 24 hours, shaking at 37°C. Fluorescence was detected with a spectral curve with excitation at 480nm and emission detected between 500nm-600nm. Relative fluorescence units (RFUs) represent raw values with media only background subtracted. TIMER clones are shown alongside the non-fluorescent *E. coli* parent strain. Data represents mean of three biological replicates. B) Gating strategy for fluorescence-activated cell sorting. FSC, SSC, and pulse width were used to select the bacterial population (within R1 and R2 gates), and cells within gate R3 were defined as red fluorescent, and were collected. C) Table summary of 4 clones selected with high red fluorescence in stationary phase *E. coli*. Clone name is shown with all mutations in the TIMER gene, and additional notes about each clone. D) Red fluorescence (560nm excitation, 610nm emission) of stationary phase (24h growth) non-fluorescent DH5αλpir *E. coli* parent strain, containing TIMER_42_ (inserted into pMMB67EH multiple cloning site), or containing *P*_*hmsT*_*∷TIMER*_*42*_ (inserted into pMMB67EH multiple cloning site). Data represents three biological replicates. Statistics: D) Kruskal-Wallis one-way ANOVA with Dunn’s post-test, *p<.05.

We hypothesized this lack of red fluorescence accumulation was because the folding of red monomeric TIMER_I161T T197A_ was still too slow, relative to the doubling times of Yptb and *E. coli*. Therefore, we used the TIMER_I161T T197A_ construct as a template for random mutagenesis, to specifically isolate TIMER variants that have red fluorescence accumulation in stationary phase cells. TIMER variants were inserted into pUC18, transformed into *E. coli*, and transformants were grown for 24h with IPTG. Non-fluorescent *E. coli* cultures were grown in parallel to define the threshold for fluorescence detection during cell sorting. Forward scatter (FSC), side scatter (SSC) and pulse width detection were used to identify individual bacterial cells, and we collected bacteria with detectable 572nm emission (gate R3) **(Fig 1B)**. Collected bacteria (approximately 200 cells) were plated for single colonies, and 192 colonies were screened for red fluorescence during growth in the presence of IPTG by plate reader. 26 clones had detectable red fluorescence in stationary phase (24h growth) compared to exponential phase (4h growth); one clone, TIMER_42_, is shown as a representative **(Fig 1A)**. Some red fluorescence was detected without IPTG induction, likely due to the high copy number of pUC18; this signal emerged between 16h and 20h growth in the presence of IPTG **(Supplemental Fig 2A**). The 26 clones were sequenced, and their TIMER sequences represented three different ‘types’, numbered based on the well number of the first observed clone. Three of the sequenced clones were the TIMER_42_ type, with a T22T synonymous mutation (ACC--> ACA) that is predicted to increase translation efficiency (29); one clone was the TIMER_48_ type, with a S69T mutation in the central helix that likely impacts chromophore maturation, since it is adjacent to chromophore residues 66-68 (11); and the other 22 clones were the TIMER_56_ type, which contain a TIMER_I161T T197A_ gene with a single adenosine insertion upstream of the start codon, likely resulting in increased transcript levels **(Fig 1C)**. Of these three clone types, TIMER_42_ and TIMER_56_ had high red fluorescent signal in stationary phase, but the signal from TIMER_48_ was dim in comparison **(Supplemental Fig 2B)**. Because of its bright signal and confirmed mutation within the TIMER gene, we selected the TIMER_42_ clone for additional experiments.

**Fig 2:**
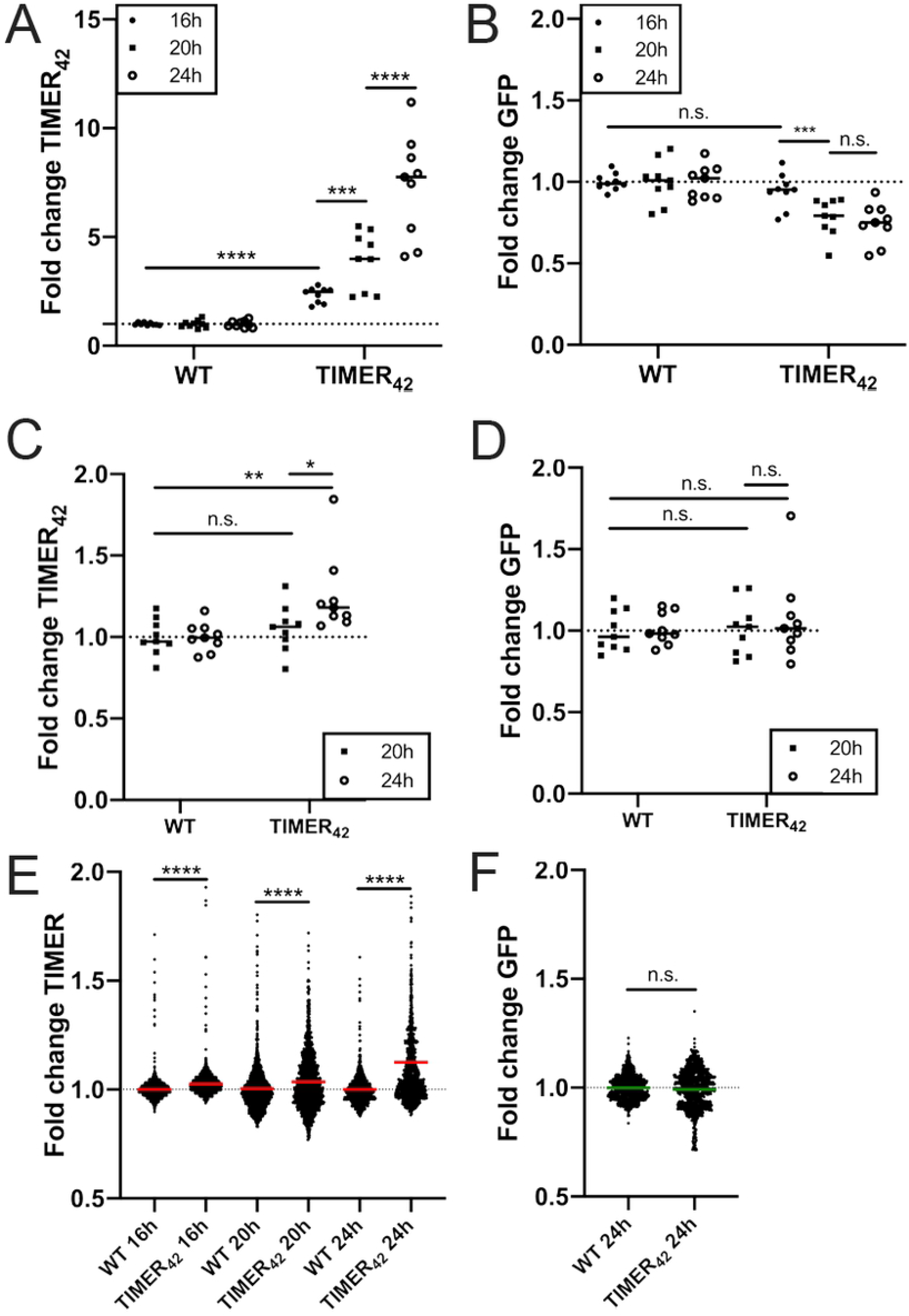
TIMER_42_ accumulates red, but not green fluorescence in stationary phase *Y. pseudotuberculosis*. WT IP2666 Yptb and *P*_*hmsT*_*∷TIMER*_*42*_ Yptb strains were grown in 96 well plates at 37°C with shaking, in the presence or absence of 1mM IPTG, for the indicated timepoints (hours: h). Fluorescence was detected by plate reader and fluorescence microscopy. A) Plate reader detection of TIMER_42_ signal accumulation with IPTG induction. Dots: biological replicates. B) Plate reader detection of green fluorescence with IPTG induction, same samples as A). Dots: biological replicates. C) Plate reader detection of TIMER_42_ signal accumulation without IPTG. Dots: biological replicates. D) Plate reader detection of green fluorescence without IPTG, same samples as C). Dots: biological replicates. E) Fluorescence microscopy to detect TIMER_42_ signal within individual bacterial cells, grown for the indicated times in the absence of IPTG. Dots: individual cells. F) Green fluorescence from bacterial cells shown in E). Fold change: signal relative to non-fluorescent WT IP2666, grown alongside TIMER_42_ strains under the same conditions and timepoints. Data represents 9 biological replicates for each condition. Dotted lines: baseline value of non-fluorescent WT IP2666 cells. Bars: mean values. Statistics: A)-D) Two-way ANOVA with Tukey’s multiple comparison test; E) Kruskal-Wallis one-way ANOVA with Dunn’s post-test; F) Mann-Whitney; ****p<.0001, ***p<.001, **p<.01, *p<.05, n.s.: not-significant.

We then sought to move TIMER_42_ into the low copy vector pMMB67EH, which is stably maintained during mouse infection (22, 26). Insertion of TIMER_42_ into the pMMB67EH multiple cloning site (MCS) did not result in detectable fluorescence despite the upstream *P*_*tac*_ promoter, so we also inserted *P*_*hmsT*_, the promoter for the diguanylate cyclase *hmsT*, upstream of TIMER_42_. Our *Y. pseudotuberculosis* strain (IP2666) naturally contains a disrupted Fur binding site in *P*_*hmsT*_, which should result in low, constitutive transcription of genes downstream of *P*_*hmsT*_ (30, 31). *P*_*hmsT*_ was sufficient to promote the accumulation of red TIMER_42_ monomers in stationary phase cells (24h growth), in the absence of IPTG induction **(Fig 1D)**.

### TIMER_42_ accumulates red, but not green fluorescence in stationary phase *Y. pseudotuberculosis*

The *P*_*hmsT*_*∷*TIMER_42_ construct was then transformed into WT *Y. pseudotuberculosis* (Yptb) to determine if TIMER_42_ signal accumulation could be detected in stationary phase Yptb. Initially, we utilized IPTG induction, and detected fluorescent signal accumulation using a plate reader. A significant increase in TIMER signal was detected after 16h (hours) of growth in the presence of IPTG **(Fig 2A)**. TIMER signal continued to increase between 20h and 24h of growth in the presence of IPTG, confirming TIMER signal accumulates in late stationary phase Yptb. We hoped to utilize green fluorescent reporters alongside our TIMER constructs, so it was important to determine if the TIMER_42_ construct would produce interfering fluorescence across green wavelengths. Despite high TIMER signal accumulation, no significant green fluorescence was detected **(Fig 2B)**. We then asked whether TIMER signal would accumulate in the absence of IPTG-induction, since we would not be inducing with IPTG in our mouse model of deep tissue infection, and found that TIMER signal could be detected after 24h growth in LB **(Fig 2C)**. We also confirmed that cells lacked green fluorescent signal at this timepoint **(Fig 2D)**.

Plate reader detection represents fluorescent values averaged across all cells within a well of a 96-well plate. To determine if there was variability in TIMER signal accumulation within individual bacterial cells in these samples, we used fluorescence microscopy to detect fluorescence at the single cell-level. There was a significant increase in the mean TIMER_42_ signal after 16h, 20h, and 24h growth relative to non-fluorescent WT cells, but this was most apparent at 24h growth **(Fig 2E)**. The observed variability across the bacterial population suggests that subsets of cells are truly slow-growing, TIMER_42+_, at each of these timepoints, while cells lacking TIMER_42_ signal accumulation may remain more metabolically active. We also confirmed that cells lacked green fluorescent signal after 24h growth, again suggesting that GFP reporters can be used alongside TIMER_42_ **(Fig 2F)**. Collectively, these data indicate that TIMER_42_ signal accumulates late in stationary phase Yptb, between 16h and 24h of culture, and that the TIMER_42_ construct lacks green fluorescent signal.

### TIMER_42_ is detected within the host spleen at 72h p.i

To quantify the relative TIMER signal accumulation within individual bacterial cells, we transformed TIMER_42_-expressing Yptb (*P*_*hmsT*_*∷*TIMER_42_) with a plasmid that constitutively-expresses GFP (22, 26). To determine when TIMER_42_ signal accumulated during bacterial growth in the spleen, we intravenously infected C57BL/6 mice with this GFP^+^ TIMER_42_ strain and harvested spleens at 48h and 72h post-inoculation (p.i.) to detect TIMER_42_ signal. TIMER_42_ signal was not detected in microcolonies at 48h p.i., but was detectable at 72h p.i. (**Fig 3A**). To determine if TIMER_42_ signal accumulated in a specific spatial location within microcolonies, we compared the ratio of TIMER_42_/GFP signal intensity at the geometric centroid and periphery of individual microcolonies (**Fig 3B**). There was not a significant difference in the relative TIMER_42_ signal at the centroid or periphery, suggesting TIMER_42_ accumulation does not occur in a specific spatial location across microcolonies. We also compared the relative TIMER_42_ signal at the centroid and periphery of each microcolony, and found a slight positive correlation in the two signals (**Fig 3C**). This shows that higher centroid signal correlates with higher periphery signal, and further suggests that on average, the TIMER_42_ signal isn’t specifically enriched at either the centroid or the periphery. However, TIMER_42_ signal accumulation also did not appear uniform across microcolonies, which could suggest that specific subsets of cells may be transitioning to a slowed growth phenotype.

**Fig 3:**
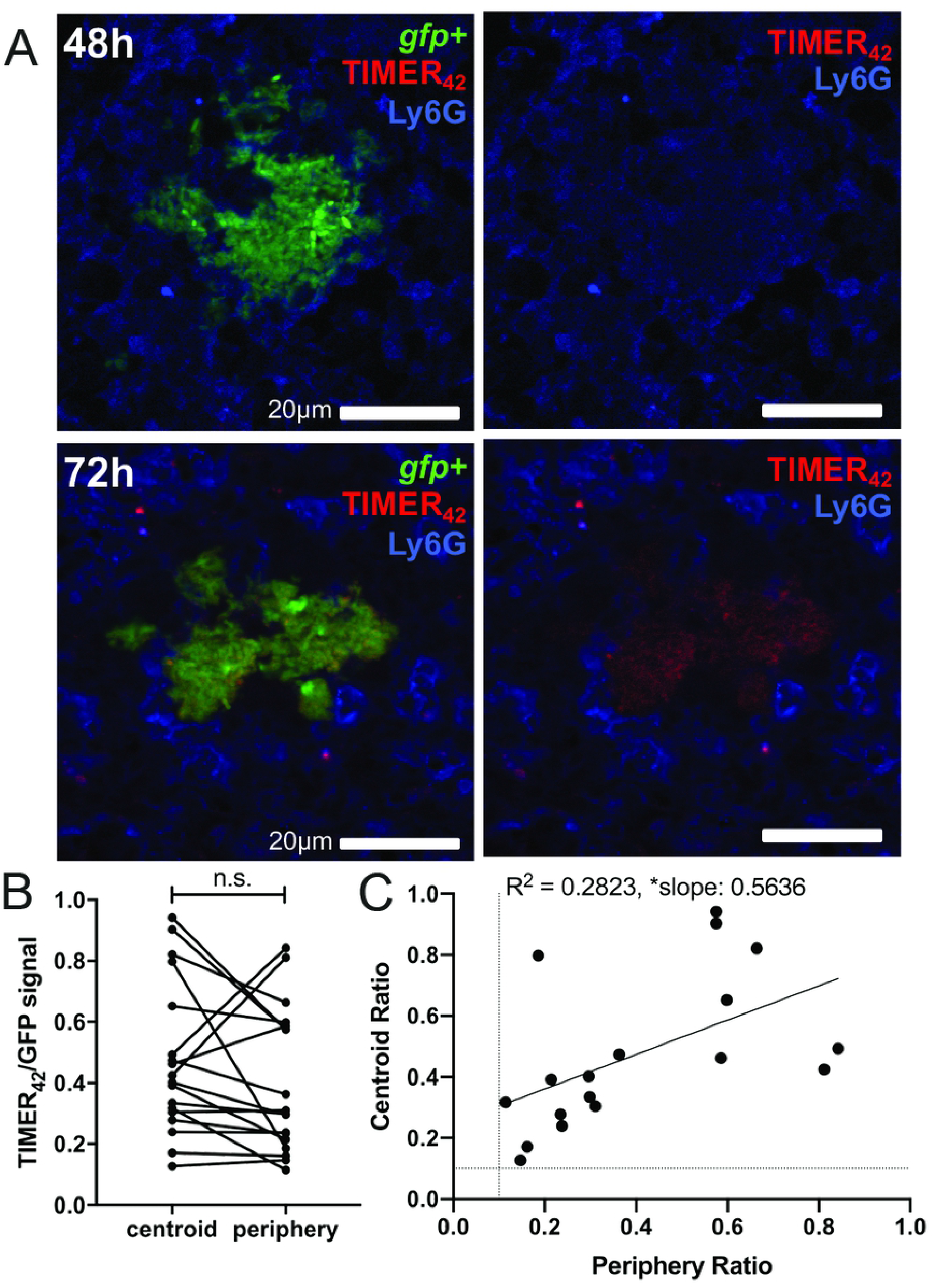
TIMER_42_ is detected within the host spleen at 72h p.i. C57BL/6 mice were infected intravenously with the TIMER_42_ (*P*_*hmsT*_*∷TIMER*_*42*_) GFP^+^ strain for the indicated timepoints post-inoculation (p.i.) A) Representative images are shown at 48h and 72h as merged images (left), and without the green channel (right). Tissues were also stained for neutrophils (Ly6G). B) TIMER_42_ signal intensity was divided by GFP signal intensity (TIMER_42_/GFP signal) at the centroid and periphery. Each dot: individual microcolony. Lines connect centroid and periphery values from the same microcolony. C) The TIMER_42_/GFP ratio at the centroid was plotted relative to the periphery value for each microcolony. Each dot: individual microcolony. Dotted lines: threshold for signal detection; only microcolonies with detectable TIMER_42_ signal were included in the analysis. Statistics: B) Wilcoxon-matched pairs; C) linear regression with R^2^ value and slope of best-fit line; *p<.05, n.s.: not significant. Significantly non-zero slope indicates values are correlated.

### TIMER_42_ signal accumulates after stationary phase markers

To determine if there are specific subsets of Yptb that exhibit TIMER_42_ accumulation and slowed growth, we compared TIMER_42_ signal to expression from several different fluorescent reporters. We began by testing TIMER_42_ alongside two different *dps* reporter constructs, which should also detect slow-growing subsets of cells. *dps* expression occurs when bacteria transition into stationary phase, and *dps* gene expression has been used extensively as a marker of slowed growth (32). However, Dps protein levels are regulated at the transcriptional, post-transcriptional, and translational levels by multiple pathways, suggesting that a *dps* transcriptional reporter alone may not be sufficient to identify slow growing cells, necessitating a tool like TIMER_42_ (33). We generated two *dps* reporters: *P*_*dps*_*∷gfp* and *P*_*dps*_*∷gfp-ssrA*, where the *dps* promoter is fused to stable and destabilized *gfp*, respectively (34, 35). This destabilized version of *gfp* has a half-life of approximately 37 minutes in culture, compared to approximately 147 minutes for stable *gfp*, indicating the destabilized GFP signal will depict recent transcriptional changes (**Supplemental Fig 3**). *dps* reporters were transformed into TIMER_42_-expressing Yptb (TIMER_42_ *P*_*dps*_*∷gfp* or TIMER_42_ *P*_*dps*_*∷gfp-ssrA*), individual colonies were inoculated into 96 well plates, and fluorescent signals were detected over time. Expression of the stable *dps* reporter was first detected after 5h growth, and remained relatively steady for about 12h, until we saw a gradual increase in signal (**Fig 4A**). In contrast, the destabilized *dps* reporter had very little fluorescence until 20h growth, which was similar to the pattern with TIMER_42_ signal accumulation in the same strain background **(Fig 4A**). TIMER_42_ signal accumulation was very similar in both strain backgrounds; only TIMER_42_ signal in TIMER_42_ *P*_*dps*_*∷gfp-ssrA* is shown to directly compare the two signals in this strain. These data suggested that the stable *P*_*dps*_*∷gfp* reporter signal accumulates prior to the stationary phase transition, and that the destabilized *P*_*dps*_*∷gfp-ssrA* reporter is more accurately detecting slow growing cells.

**Fig 4:**
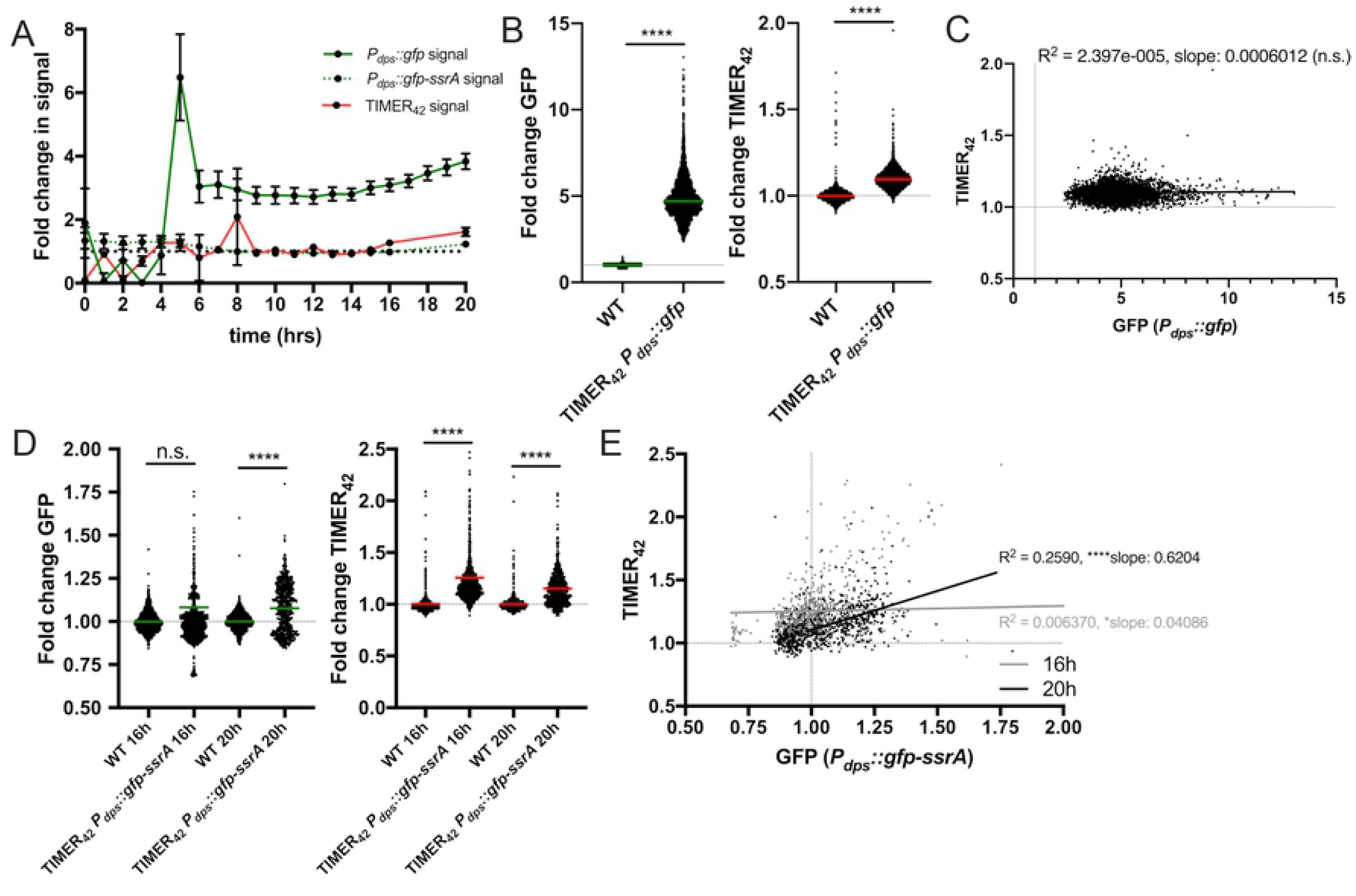
TIMER_42_ signal accumulates after stationary phase markers. *P*_*dps*_*∷gfp* TIMER_42_, *P*_*dps*_*∷gfp-ssrA* TIMER_42_ and WT strains were grown in 96 well plates at 37°C under standing conditions for the indicated timepoints, and fluorescence was detected by plate reader and fluorescence microscopy. A) Timecourse of fluorescent signal detection by plate reader, shown as fold change relative to non-fluorescent WT cells. TIMER_42_ data shown from the *P*_*dps*_*∷gfp-ssrA* TIMER_42_ strain, to compare with *P*_*dps*_*∷gfp-ssrA* reporter signal; very similar results were obtained with TIMER_42_ signal from the *P*_*dps*_*∷gfp* TIMER_42_ strain. Plate reader data represents 15 biological replicates for each strain. B) Fold change GFP and TIMER_42_ within individual bacteria quantified by fluorescence microscopy. Indicated strains were grown 16 hours. C) Correlation plot of fold change in TIMER_42_ and GFP signals from bacteria in B). Dots: individual cells. D) Fold change GFP and TIMER_42_ within individual bacteria quantified by fluorescence microscopy. Indicated strains were grown 16 or 20 hours. E) Correlation plot of fold change in TIMER_42_ and GFP signals from bacteria in D). Dots: individual cells. Dotted lines: baseline value of non-fluorescent WT cells. Microscopy data represents 4 biological replicates for each strain and condition. Statistics: B) Mann-Whitney; C) & E) linear regression with R^2^ value and slope of best-fit line; D) Kruskal-Wallis one-way ANOVA with Dunn’s post-test; ****p<.0001, *p<.05, n.s.: not-significant. Significantly non-zero slope indicates values are correlated.

There was a significant increase in both *P*_*dps*_*∷gfp* and TIMER_42_ signal after 16h growth, as detected by fluorescence microscopy (**Fig 4B**), however there was no correlation between these two signals within individual bacteria (**Fig 4C**), likely due to the early accumulation of stable GFP. However, with the TIMER_42_ *P*_*dps*_*∷gfp-ssrA* strain, significant GFP signal was not detected until 20h growth by fluorescence microscopy, while TIMER_42_ was detected at both 16h and 20h growth (**Fig 4D**). There was a significant correlation between *P*_*dps*_*∷gfp-ssrA* and TIMER_42_ signal at 20h growth, suggesting these reporters are detecting the same subsets of cells, and are behaving similarly at this late timepoint (**Fig 4E**). For additional information about statistical analyses for correlations, see Materials and Methods below. Collectively, this indicates that both *dps* and TIMER_42_ can be used to identify slow growing cells. However, *dps* signal detection is heavily dependent on the stability of GFP, and an early increase in *gfp* transcripts at the beginning of the transition into stationary phase can lead to prolonged signal detection. TIMER_42_ signal accumulation is detected well after this point, and likely means that TIMER_42_ can be used to detect cells much later in stationary phase.

### TIMER_42_ signal accumulation correlates with *hmp* expression during exposure to acidified nitrite

We then sought to determine if the response to reactive nitrogen species (RNS) is sufficient to slow the growth of cells and promote TIMER signal accumulation, since we observe bacterial responses to this stress during growth in the spleen (22, 26). To determine if RNS were sufficient to promote TIMER_42_ signal accumulation, we first used the acidified nitrite method of generating nitric oxide (NO), where NaNO_2_ is added to acidified LB (pH 5.5) (22, 26, 36). The 2.5mM NaNO_2_ concentration was known to inhibit growth and promote high levels of *hmp* expression, whereas 0.25mM represents a sub-inhibitory concentration (26). *hmp* and TIMER_42_ fluorescence were detected within individual cells by fluorescence microscopy. At 2h post-treatment, all concentrations of NaNO_2_ promoted *hmp* expression, and there was significantly higher *hmp* expression with the highest concentration, 2.5mM (**Fig 5A**). TIMER_42_ signal accumulation was also detected following exposure to 1mM and 2.5mM NaNO_2_, but TIMER_42_ signal was highest with 1mM NaNO_2_ (**Fig 5B**). TIMER_42_ and *hmp* signals were correlated during 1mM NaNO_2_ treatment, but 2.5mM NaNO_2_ resulted in a significant negative correlation, suggesting this high concentration was impacting bacterial viability due to high levels of stress (**Fig 5C**). At 4h post-treatment, the highest *hmp* reporter expression was detected with the 1mM intermediate dose, and the *hmp* signal had decreased with the 2.5mM dose, again suggestive of high levels of stress (**Fig 5D**). TIMER_42_ signal was detected with all three doses, and was highest with 2.5mM (**Fig 5E**). However, there was an interesting divergence in the 1mM population, where both TIMER42^+^ TIMER42^-^ cells were observed, suggesting that some cells may have recovered from stress and resumed growth resulting in a loss of TIMER_42_ signal (**Fig 5E**). We observed a significant positive correlation between TIMER_42_ and *hmp* signals with the low 0.25mM and intermediate 1mM doses at this timepoint, but no correlation with the highest 2.5mM dose (**Fig 5F**). Collectively, these results show that acidified nitrite is sufficient to promote TIMER signal accumulation, and suggest TIMER_42_ can be used to detect growth-arrested cells.

**Fig 5:**
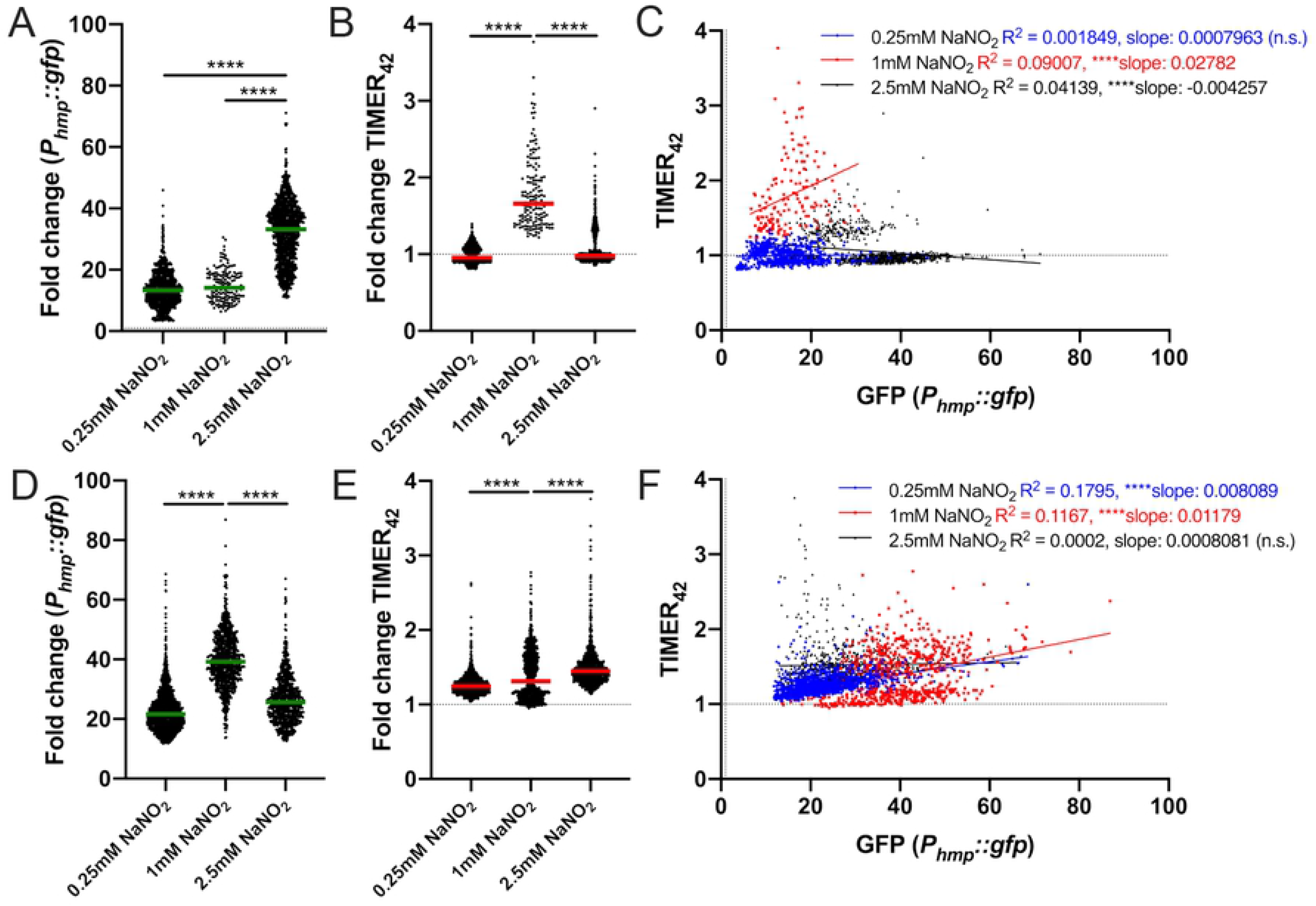
TIMER signal accumulation correlates with *hmp* expression during exposure to acidified nitrite. *P*_*hmp*_*∷gfp* TIMER_42_ and WT strains were grown in LB pH5.5 in 96 well plates at 37°C with shaking, with the indicated concentrations of NaNO_2_, for either 2h (panels A-C) or 4h (panels D-F). Fluorescence of individual bacteria was detected by fluorescence microscopy. Fold change in signal is shown relative to non-fluorescent WT bacteria treated under the same conditions (dotted lines). A) GFP fluorescence after 2h treatment. B) TIMER_42_ fluorescence after 2h treatment.. C) Correlation plot of fold change in TIMER_42_ and GFP signals from bacteria in A) and B). D) GFP fluorescence after 4h treatment. E) TIMER_42_ fluorescence after 4h treatment. F) Correlation plot of fold change in TIMER_42_ and GFP signals from bacteria in D) and E). Dots: individual cells. Dotted lines: baseline value of non-fluorescent WT cells. Microscopy data represents 4 biological replicates for each strain and condition. Statistics: A), B), D), E): Kruskal-Wallis one-way ANOVA with Dunn’s post-test; C) & F): linear regression with R^2^ value and slope of best-fit line; ****p<.0001, *p<.05, n.s.: not-significant. Significantly non-zero slope indicates values are correlated.

### NO stress is sufficient to promote TIMER_42_ signal accumulation

The results with acidified nitrite suggest that RNS are sufficient to promote TIMER_42_ signal accumulation, but to truly show nitric oxide (NO) exposure is sufficient, we performed experiments with the DETA-NONOate NO donor compound. Cultures of the TIMER_42_ *P*_*hmp*_*∷gfp* and TIMER_42_ *P*_*hmp*_*∷gfp-ssrA* strains were exposed to 2.5mM DETA-NONOate for 4h, and fluorescence microscopy was used to detect fluorescent signals within individual bacterial cells. The 2.5mM dose was chosen for these experiments based on our previous studies, which showed this dose is sufficient to promote *hmp* reporter expression (22, 26, 37). Treatment with the NO donor compound resulted in high levels of *hmp* reporter expression with both strains (**Fig 6A**), and TIMER_42_ signal accumulation (**Fig 6B**). TIMER_42_ and *hmp* signal were significantly correlated for both the stable reporter strain (**Fig 6C**) and the unstable reporter strain (**Fig 6D**), showing that NO is sufficient to promote TIMER signal accumulation.

**Fig 6:**
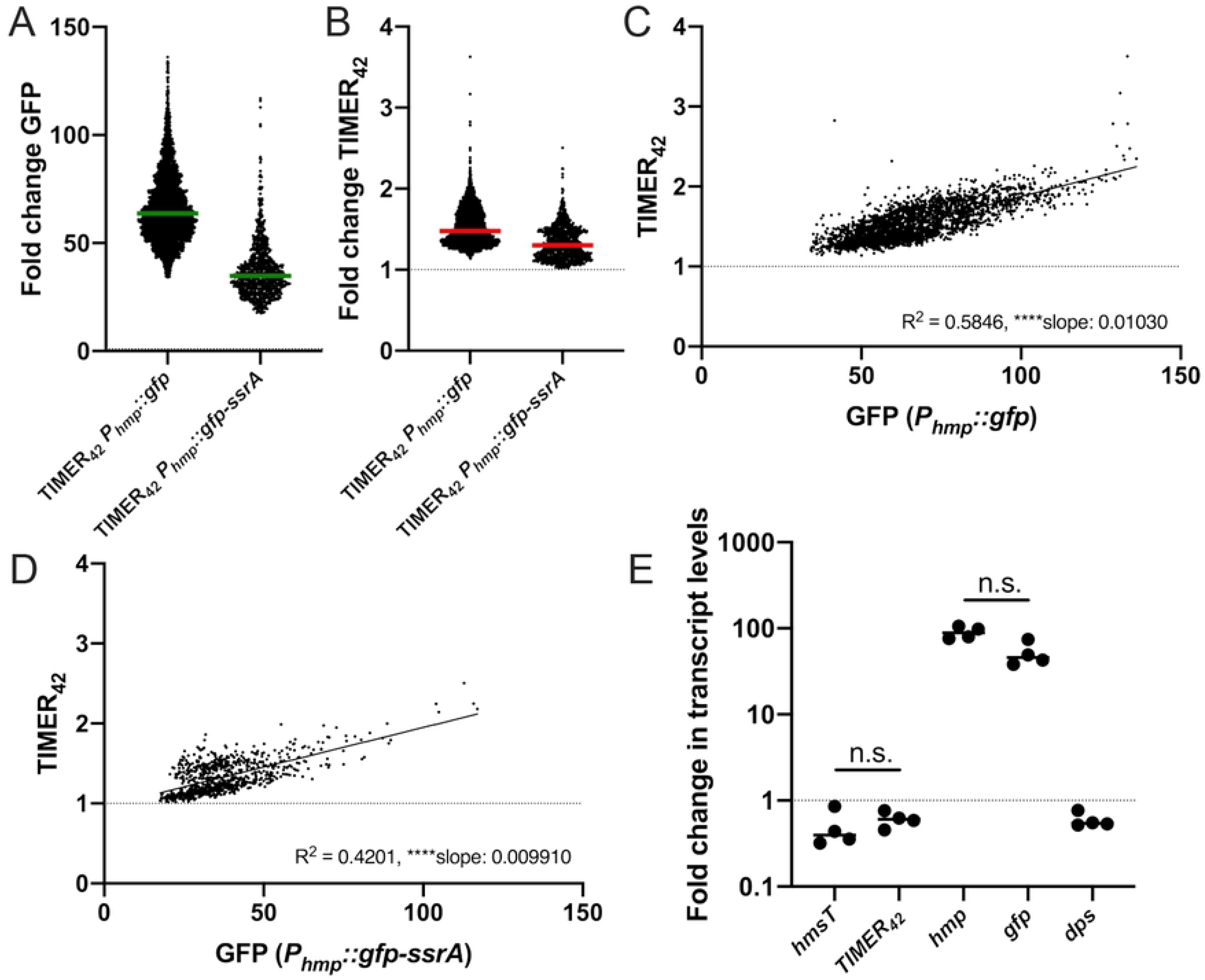
NO stress is sufficient to promote TIMER_42_ signal accumulation. *P*_*hmp*_*∷gfp* TIMER_42_, *P*_*hmp*_*∷gfp-ssrA* TIMER_42_, and WT strains were grown in LB in 96 well plates at 37°C with shaking, with 2.5mM DETA-NONOate for 4h. A) GFP fluorescence of individual bacteria of the indicated strains was detected by fluorescence microscopy. Fold change in signal is shown relative to non-fluorescent WT bacteria treated under the same conditions (dotted lines). B) TIMER_42_ fluorescence of bacteria shown in A). C) Correlation plot of fold change in TIMER_42_ and GFP signals from bacteria in A) and B), *P*_*hmp*_*∷gfp* TIMER_42_ strain. D) Correlation plot of fold change in TIMER_42_ and GFP signals from bacteria in A) and B), *P*_*hmp*_*∷gfp-ssrA* TIMER_42_ strain. Dots: individual cells. Dotted lines: baseline value of non-fluorescent WT cells. Microscopy data represents 5 biological replicates for each strain and condition. E) qRT-PCR detection of the indicated transcripts (x-axis) in *P*_*hmp*_*∷gfp* TIMER_42_ bacteria during DETA-NONOate exposure, using conditions described above. Dots: individual replicates. Fold change in transcript levels relative to untreated WT cells. Four biological replicates are shown. Statistics: C)-D): linear regression with R^2^ value and slope of best-fit line; E): Kruskal-Wallis one-way ANOVA with Dunn’s post-test; ****p<.0001, n.s.: not-significant. Significantly non-zero slope indicates values are correlated.

We also performed qRT-PCR with the TIMER_42_ *P*_*hmp*_*∷gfp* strain to confirm that TIMER_42_ signal accumulation in response to NO was not due to an increase in TIMER_42_ transcript levels, and to verify that *hmsT* was not increasing in transcript levels during NO exposure, since this promoter was used to drive TIMER_42_ expression. TIMER_42_ *P*_*hmp*_*∷gfp* cells were exposed to 2.5mM DETA-NONOate for 4h and transcripts were detected by qPCR. TIMER_42_ and *hmsT* transcript levels were not impacted by NO, and high levels of both *hmp* and *gfp* transcripts were detected, consistent with TIMER_42_ accumulation as a result of protein folding, and *gfp* expression as a result of transcription downstream of *P*_*hmp*_ (**Fig 6E**). We also quantified *dps* transcript levels to determine if these increased in response to NO, and found there was no change in *dps* expression in response to NO (**Fig 6E**). These results provide additional insight into the mechanism of NO-mediated stress, and suggest the response to NO is sufficient to arrest cell division, but not promote a stationary phase transition. This is consistent with published results showing NO disrupts cell division by arresting assembly of the FtsZ cytokinetic ring (38, 39).

### TIMER_42_ signal correlates with *dps* expression *in vivo*

We then asked whether there was a correlation between TIMER_42_ signal accumulation and either *dps* or *hmp* reporter expression during growth within host tissues. C57BL/6 mice were inoculated intravenously with either the TIMER_42_ *P*_*dps*_*∷gfp* or TIMER_42_ *P*_*hmp*_*∷gfp* strains, and spleens were harvested at 72h p.i. to visualize reporter expression by fluorescence microscopy. At 72h p.i., some microcolonies had detectable TIMER_42_ signal accumulation (TIMER_42+_), while other microcolonies lacked TIMER signal (TIMER_42-_). This was also observed in initial experiments with the GFP^+^ TIMER_42_ strain, and only TIMER_42+_ microcolonies were analyzed in Fig 3. We hypothesized that larger microcolonies may run out of nutrients, which then slows growth, and results in TIMER_42_ signal accumulation. We compared the areas of microcolonies that had or lacked TIMER_42_ signal, and found that generally the TIMER_42+_ microcolonies were indeed larger (**Fig 7A**).

**Fig 7:**
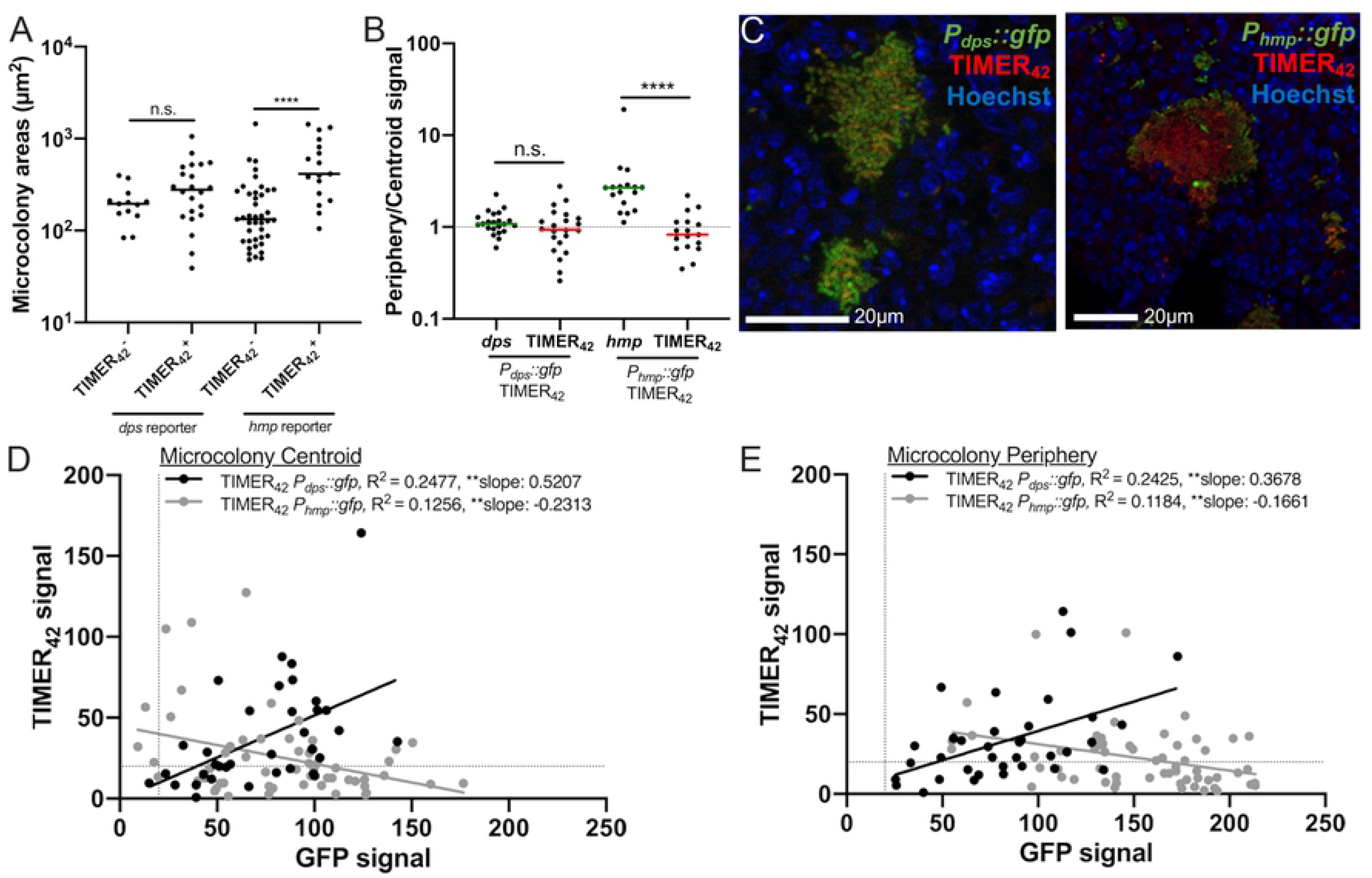
TIMER_42_ signal correlates with *dps* expression *in vivo*. C57BL/6 mice were infected intravenously with the TIMER_42_ (*P*_*hmsT*_*∷TIMER*_*42*_) *P*_*dps*_*∷gfp* or TIMER_42_ *P*_*hmp*_*∷gfp* strain for 72h, and spleens were harvested for fluorescence microscopy. A) Areas of individual microcolonies, separated based on whether TIMER_42_ signal was detectable above background (TIMER 42^+^) or undetectable (TIMER 42 ^-^). Data with the TIMER42 *Pdps ∷gfp* strain (*dps* reporter) and TIMER_42_ *P*_*hmp*_*∷gfp* strain (*hmp* reporter) is shown. B) Peripheral fluorescent signal intensity was divided by centroid fluorescent intensity for each reporter (x-axis). At each spatial location, *dps* or *hmp* reporter signal was detected alongside TIMER_42_ signal, to compare their signal intensities directly. Data represent microcolonies defined as TIMER_42+_. Dotted line: value of 1, where signal is equivalent at the centroid and periphery (no spatial patterning). C) Representative images. D) TIMER_42_ signal was plotted relative to either *dps* or *hmp* reporter signal to determine if the signal intensities were correlated at the centroid of microcolonies. All microcolonies detected within tissues are included (those defined as TIMER_42+_ and TIMER_42-_). D) TIMER_42_ signal was plotted relative to either *dps* or *hmp* reporter signal to determine if the signal intensities were correlated at the periphery of microcolonies. Each dot: individual microcolony. Dotted lines: threshold for signal detection. Statistics: A)-B) Wilcoxon-matched pairs; D)-E) linear regression with R^2^ value and slope of best-fit line; **p<.01, *p<.05, n.s.: not significant. Significantly non-zero slope indicates values are correlated.

We then asked whether *dps* or *hmp* reporter signal correlated with TIMER_42_ signal accumulation within these tissues. We first compared the ratio of *dps* and TIMER_42_ signal at the periphery and centroid of microcolonies and found the signal intensity of each of these reporters looked very similar (**Fig 7B**). In comparison with TIMER_42_, *hmp* reporter signal was significantly enriched at the periphery (**Fig 7B**). This *hmp* reporter signal pattern was consistent with our previously published results (22, 26, 37) but suggested that NO stress was not significantly slowing bacterial growth within host tissues, based on TIMER_42_ signal accumulation in some, but not all, peripheral cells. Representative images indicate there was substantial overlap between *dps* and TIMER_42_ signals, and that *hmp* and TIMER_42_ are detecting distinct subsets of bacteria (**Fig 7C**). To further analyze the relationship between these signals, we compared the TIMER_42_ and GFP signals at microcolony centroids (**Fig 7D**) and microcolony peripheries (**Fig 7E**). For each individual microcolony spatial location, the relative TIMER_42_ and GFP signals are plotted on the Y- and X-axis respectively, to determine if there is a linkage between the two values by linear regression (see Materials and Methods below). At both the centroid and periphery, we found a significant positive correlation between TIMER_42_ and *dps* reporter signal, however we found a significant negative correlation between TIMER_42_ and *hmp* reporter signal. This supports the conclusion that *dps* and TIMER_42_ are marking the same subsets of bacteria within host tissues, but suggests that *hmp* and TIMER_42_ are detecting distinct subsets. This suggests that at 72h p.i., there is a subset of slow-growing Yptb at the interior of microcolonies, and also suggests that NO stress is not sufficient to cause prolonged growth arrest within host tissues.

### TIMER_42_ signal dissipates after initial responses to NO, suggesting recovery occurs *in vivo*

We were surprised to find that NO stress did not correlate with TIMER_42_ signal accumulation within host tissues, and hypothesized this could be due to temporal differences in when bacteria were exposed to NO and when tissues were collected. We believe that bacteria respond to NO much earlier during infection (approximately 48h p.i., (22, 28)), and it was possible that bacteria were no longer actively responding to stress. We returned to the acidified nitrite method of introducing NO stress, since this likely results in an initial burst of NO, and then slower NO generation (36). We exposed the TIMER_42_ *P*_*hmp*_*∷gfp* cells to the indicated concentrations of NaNO_2_ in LB pH5.5, and measured absorbance (A_600nm_) and reporter signal by fluorescence, in a plate reader over time. Consistent with the predicted initial burst of NO release, we initially observed growth arrest after treatment followed by slowed growth (**Fig 8A**). There was a significant increase in *hmp* reporter signal after 1h treatment that continued to increase over the course of 8h (**Fig 8B**). However, TIMER_42_ signal accumulation was consistent with an initial arrest of the cell population followed by recovery and resumed growth, based on significantly higher TIMER_42_ signal accumulation with 1mM treatment relative to 0.25mM treatment at 2h and 4h post-treatment, and a return to baseline levels 6h after the initial exposure to acidified nitrite (**Fig 8C**). Consistent with this TIMER_42_ result, growth also appeared to resume at these timepoints in cultures, based on absorbance (**Fig 8A**).

**Fig 8:**
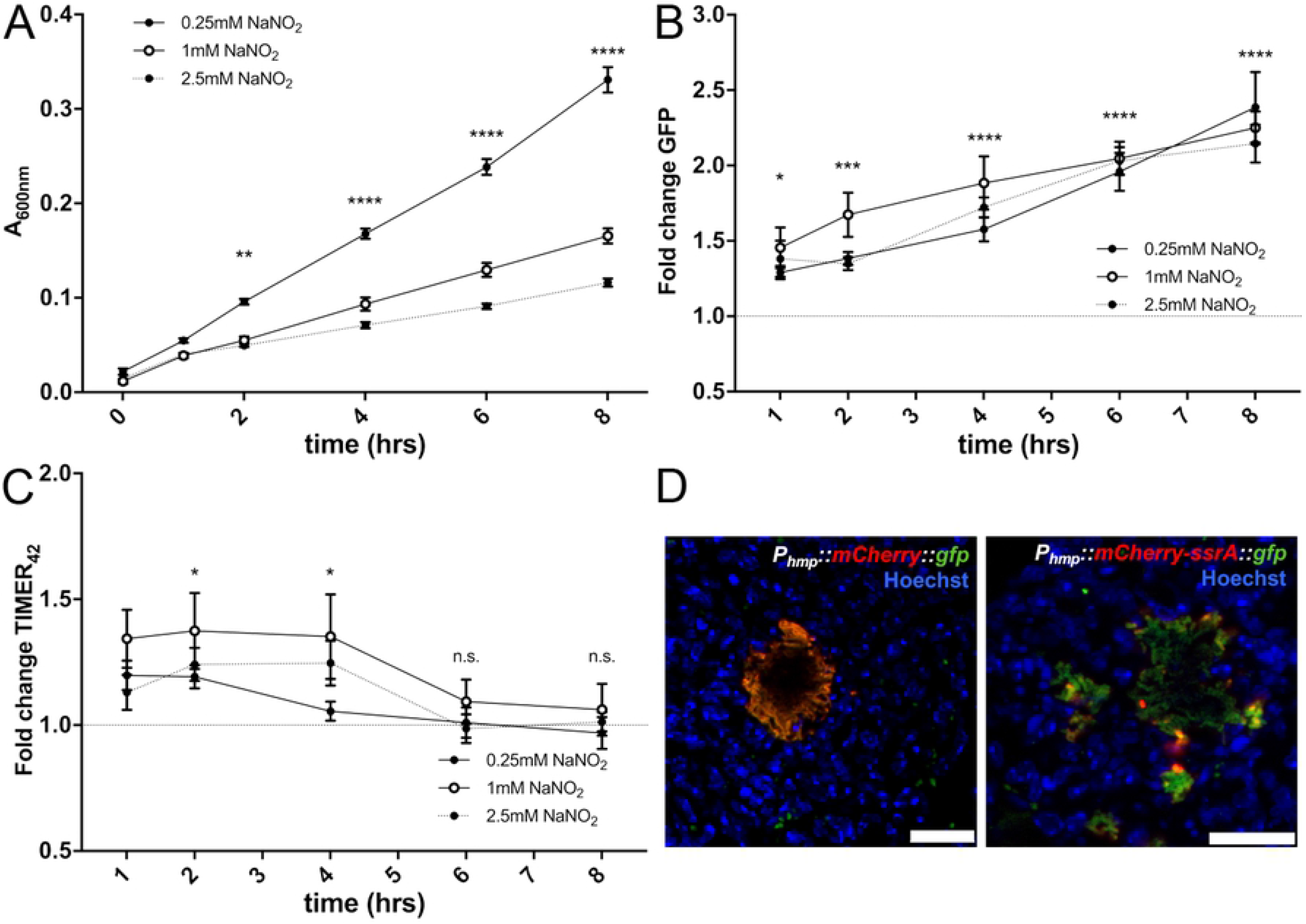
TIMER signal dissipates after initial responses to NO, suggesting recovery occurs *in vivo*. *P*_*hmp*_*∷gfp* TIMER_42_ and WT strains were grown in LB pH5.5 in 96 well plates at 37°C with shaking, with the indicated concentrations of NaNO_2_, for the indicated timepoints. A) Optical density of the *P*_*hmp*_*∷gfp* TIMER_42_ strain was measured by absorbance (A_600nm_) over time (hours, hrs), in a plate reader. B) *hmp* reporter expression (GFP signal) was detected over time by plate reader. Fold change is relative to WT non-fluorescent samples grown under the same conditions. C) TIMER_42_ signal was detected over time by plate reader. Fold change is relative to WT non-fluorescent samples grown under the same conditions. D) C57BL/6 mice were infected intravenously with strains containing either the *P*_*hmp*_*∷mCherry∷gfp* or *P*_*hmp*_*∷mCherry-ssrA∷gfp* reporter constructs, and spleens were harvested for fluorescence microscopy at 72h p.i. Representative images are shown. Nine biological replicates are shown for plate reader data with mean and standard deviation. Statistics: A)-C): Two-way ANOVA with Tukey’s multiple comparison test; ****p<.0001, ***p<.001, **p<.01, *p<.05, n.s.: not-significant. Statistics shown represent: A) and C) pair-wise comparisons between 0.25mM and 1mM treatment conditions; B) comparison between 1mM treatment and untreated cells

NO exposure has recently been shown to arrest bacterial cell division through disruption of cytokinesis machinery, and so it was expected that TIMER_42_ signal would accumulate in growth-arrested, NO-exposed, bacteria (39). The lack of correlation between the two signals during growth in tissue was unexpected, and suggested that NO stress may have occurred much earlier during infection than the 72h timepoint. To test this hypothesis, we generated constructs that contained combinations of stable and unstable fluorescent protein genes with expression driven by the *P*_*hmp*_ promoter. *P*_*hmp*_*∷mCherry∷gfp* was a control construct generated with stable mCherry and stable GFP proteins, and we confirmed these signals overlapped during growth in tissue (**Fig 8D**). The *P*_*hmp*_*∷mCherry-ssrA∷gfp* construct contained destabilized mCherry (28) and stable gfp, to allow us to detect bacteria that had responded to NO as some point during infection (GFP^+^) compared to bacteria that were actively responding to NO when the tissue was harvested and fixed (mCherry^+^). Using this reporter, we found very few bacterial cells were actively responding to NO when tissue was harvested (**Fig 8D**), suggesting that other cells may have responded to NO, recovered, and resumed growth, explaining why they no longer exhibited TIMER_42_ signal accumulation. Instead, TIMER_42_ signal accumulation is clearly marking an interior subset of bacteria, that are likely experiencing nutrient limitation at this late timepoint post-inoculation.

## Discussion

In this study, we sought to modify TIMER to detect slow-growing bacteria within host tissues, with a goal of generating a tool to enable future studies of slow-growing bacterial populations. The TIMER_42_ variant has been tested in *E. coli* and *Y. pseudotuberculosis*; the signal accumulates late in stationary phase in both of these quickly-replicating Gram-negative organisms. TIMER_42_ is clearly visible within host tissues, and is also visible within cultured individual cells. However, the fold change quantification we used here indicate that the signal is sometimes dim, so non-fluorescent cells need to be included as a control for all assays to define TIMER42^+^ cells. TIMER42 could potentially be used directly in other Gram-negative species, or used as a template for random mutagenesis to select an optimized TIMER variant in another system. Additional rounds of random mutagenesis and sorting could also be used to select for a brighter TIMER variant, and again, this may be needed to apply TIMER_42_ to other bacterial systems. We believe the TIMER signal generated with this variant is sufficient to isolate TIMER_42_-expressing cells from host tissues, and plan to analyze these cells in future experiments to better understand the pathways promoting slowed bacterial growth.

TIMER signal accumulation requires folding prior to cell division events, and so additional variants that fold more quickly could lead to increased signal accumulation. We have shown here that TIMER_42_ signal accumulation in response to NO is likely due to protein folding during an arrest in cell division, and not changes in transcript level, based on similar *TIMER*_*42*_ transcript levels after NO donor addition (**Fig 6C**). However, it does remain somewhat unclear why the T22T synonymous mutation (ACC→ACA) is sufficient to promote TIMER_42_ signal accumulation in stationary phase cells. We initially thought this change may result in use of a more favorable threonine codon (ACA), but the original threonine codon (ACC) is used more frequently across the *Y. pseudotuberculosis* genome (40). Instead, we believe the ACA codon may increase the stability of TIMER mRNA, leading to increased levels of translation of the TIMER_42_ variant relative to TIMER_I161T T197A_, based on the predicted influence of an ACC→ ACA change in *E. coli* (29). These results in *E. coli* suggest that protein levels correlate better with mRNA concentrations than genomic codon-usage frequency, and that specific codons (including ACA) at the 5’ end of mRNA molecules could have a significant impact on mRNA stability and subsequent protein levels. Based on the predictions that codons ending in ‘A’ are generally more favorable and lead to increased protein expression, we postulate that this likely underlies the increased signal accumulation with TIMER_42_.

Recently, we have shown that NO-exposed Hmp^+^ cells preferentially survive doxycycline treatment in our mouse model of infection, which suggested the Hmp^+^ cells may represent a slow growing subpopulation of bacteria (28). Our *in vitro* results were consistent with this, where exposure to NO was sufficient for TIMER_42_ signal accumulation. Our results are also consistent with a recently published study which showed that NO causes a collapse of proton motive force (PMF) and collapse of the divisome complex, thus resulting in cell division arrest (39). However, within host tissues, slow-growing bacterial cells with TIMER_42_ signal accumulation were largely Hmp^-^, and instead expressed the stationary phase marker, Dps. We believe this surprising result may be a reflection of the transient bacterial exposure to NO during infection. Consistent with this idea, we found that only a few bacterial cells express the destabilized *hmp* reporter at 72h p.i. **(Fig 8D**). The half-life of this destabilized mCherry protein is approximately 35 minutes in culture (28), and assuming the turnover rate is similar within host tissues, this marks a very recent exposure to NO. This mosaic of expression suggests that while many cumulative cells were exposed to NO and responded with *hmp* expression, this process is transient and actively occurring in only a few cells at a time.

Following transient exposure to NO, bacteria need to repair NO-mediated damage to survive and resume replication (38, 41). We also know repair of Fe-S clusters promotes *Y. pseudotuberculosis* survival after NO exposure, and likely promotes continued bacterial growth after stress (26). What remains unclear is why Hmp^+^ cells have decreased antibiotic susceptibility, since this does not appear to be linked to slowed cell division rates. Recently, it has been shown that slowed metabolic activity is a better correlate for decreased antibiotic susceptibility (1), which could mean that Hmp^+^ cells have decreased metabolic activity even after cell division has resumed. These results could also suggest that Hmp^+^ cells may change in another way that promotes a tolerance phenotype, possibly by upregulation or downregulation of specific genes or pathways, or utilization of less energetically favorable metabolic pathways, that again lead to decreased metabolic rates. It would also be interesting to visualize the impact of antibiotic treatment on TIMER-expressing populations, to see if TIMER^+^ *Yersinia* also have decreased antibiotic susceptibility, as you would predict based on results with *Salmonella* (15). Consistent with this idea, doxycycline treatment at 72h p.i. with *Y. pseudotuberculosis* did not significantly prolong mouse survival, suggesting the TIMER_42+_ cells may be very tolerant to antibiotics (28).

The *in vitro* studies we performed here were used to better characterize the conditions that lead to TIMER_42_ signal accumulation, and compare TIMER_42_ signal to other stress reporters. The results we obtained with NO were expected, based on the role of NO in causing cell division arrest. However, the results comparing TIMER_42_ with the *dps* reporter were less clear, especially as we compared TIMER_42_ and *dps* reporters in culture to the results we observed within host tissues. Based on the results *in vitro*, we believe that *dps* transcript levels begin to accumulate early in the transition into post-exponential and stationary phase. This early increase in transcripts leads to an early accumulation in the stable *dps* reporter signal, prior to the point where growth actually slows. *In vitro* results with the unstable *dps* reporter suggest there is a second wave of *dps* transcription late in stationary phase that correlates with slowed growth and TIMER_42_ signal accumulation. Our *in vivo* results instead show overlap in the stable *dps* reporter signal and TIMER_42_ signal; detection of both signals together lends confidence to the idea that these cells represent a slow-growing subset. The unstable *dps* reporter unfortunately could not be detected within host tissues, likely because the signal was so dim with this reporter construct in general (**Fig 4D**), and because the spleen has relatively high background green fluorescence. Dps protein was also enriched in TIMER^+^ *Salmonella* populations (15), again supporting the idea that the Dps^+^ TIMER42^+^ cells we detected within host tissues are truly slow-growing. Dps regulation is complex, and expression can be triggered by exposure to ROS and iron availability, in addition to RpoS-dependent stationary phase transitions and general nutrient deprivation (42-44). TIMER_42_, in contrast, will mark slowed growth or growth arrest and could be used to isolate slow-growing bacteria to better understand what stressors are promoting slowed growth. The combination of these two reporters together could be particularly informative in future experiments to determine the mechanisms underlying the formation of slow-growing subsets of bacteria within host tissues.

## Materials and Methods

### Bacterial strains & growth conditions

The WT *Y. pseudotuberculosis* strain, IP2666, was used throughout (22, 25). For all mouse infection experiments, bacteria were grown overnight (16h) to post-exponential phase in 2xYT broth (LB, with 2x yeast extract and tryptone) at 26°C with rotation. For *in vitro* experiments with *hmp* and TIMER reporters, bacteria were grown overnight (16h) to post-exponential phase in LB prior to experimental set-up. Overnight cultures of *E. coli* were grown in LB with 1mM IPTG at 37°C with rotation for 24h prior to cell sorting. 1mM IPTG was used for induction throughout these experiments.

### Site-directed and random mutagenesis

Site-directed mutagenesis was used to generate the TIMER_I161T T197A_ variant from WT TIMER. Mutations were made sequentially by amplifying across the gene with primers containing the desired mutations. Briefly, TIMER was amplified in two pieces, with the mutation incorporated at the 3’ end of the upstream piece, and 5’ end of the downstream piece; overlap extension was used to join these homologous fragments (22). We generated TIMER_I161T_ first, then used this as a template to generate TIMER_I161T T197A_. For random mutagenesis, the TIMER_I161T T197A_ gene was amplified by PCR to use as a template for random mutagenesis PCR reactions. TIMER_I161T T197A_ was then amplified using the Agilent Gene Morph II kit for a single round of random mutagenesis and PCR products were purified (QIAGEN). These products were used as template DNA in an additional PCR reaction, where products were amplified to add restriction enzyme sites onto the 5’ and 3’ ends (10 cycles). PCR products were purified again before restriction digestion (EcoRI, BamHI), ligation into the lacZ gene of pUC18, and electroporation into XL1-BLUE *E. coli*. Transformants were plated onto IPTG/X-gal LB plates to ensure the majority of colonies contained an insert, based on white colony color. Transformants were then grown in culture +IPTG (1mM) for 24h, bacteria were pelleted, resuspended in PBS, and placed on ice during fluorescence-activated cell sorting (MoFlo XDP cell sorter, Beckman Coulter). Bacteria with red fluorescence (approximately 200 cells) were collected and plated onto LB/Amp for single colonies. 192 colonies (two 96 well plates) were screened for red fluorescence after 24h growth in 200µl LB +IPTG at 37° C with shaking. 26 colonies had clear red fluorescent signal, and pUC18 plasmids were isolated by miniprep (QIAGEN) from these colonies for sequencing.

### Generation of reporter strains

Two of the fluorescent constructs in this study were previously described: the constitutive GFP and *P*_*hmp*_*∷gfp* (GFPmut3) (22, 24). The final TIMER_42_ plasmid was constructed by amplifying the *hmsT* promoter, fusing this immediately upstream of TIMER_42_, and inserting this into pMMB67EH. The destabilized *gfp* was constructed by fusing an 11 amino acid *ssrA* tag, with AAV terminal amino acids, between the last coding amino acid and the stop codon (34, 35). This was inserted downstream of the *P*_*tac*_ IPTG inducible promoter in pMMB207Δ267 (45) to quantify GFP half-life following IPTG induction. A stable *gfp* construct was generated in parallel as a control (**Supplemental Fig 3**). The *P*_*dps*_*∷gfp* and *P*_*dps*_*∷gfp-ssrA* reporters were generated by amplifying the *dps* promoter, fusing this to either stable or destabilized *gfp* by overlap extension PCR, and inserting this into pACYC184 (22). The *P*_*hmp*_*∷gfp-ssrA* reporter was generated by fusing the *hmp* promoter to destabilized *gfp*, and was inserted into pACYC184 (22). The tandem *hmp* reporter constructs were generated by first fusing either stable or destabilized mCherry to stable *gfp*; these tandems were inserted into pMMB207Δ267 first to quantify mCherry half-life (28). The *hmp* promoter was then fused to each tandem, and these fragments were inserted into pACYC184. All plasmid transformations into *Y. pseudotuberculosis* were performed using a previously published protocol (46).

### TIMER in vitro experimental design

Individual colonies were inoculated into 200µl LB within individual wells of 96-well, black-walled, clear-bottom plates (Greiner). Surrounding wells were filled with 200µl PBS to prevent evaporation during growth. Plates were covered with aluminum foil to prevent light exposure, and were incubated at 37°C with shaking for the indicated timepoints. At individual time points fluorescence and absorbance were measured using a plate reader (see below) and aliquoted for fluorescence microscopy. For experiments with *dps* reporter strains, plates were incubated without shaking at 37°C. For experiments with acidified nitrite, overnight cultures were diluted 1:50 into 200µl fresh LB pH5.5, in the presence or absence of the indicated concentrations of NaNO_2_, within wells. For experiments with DETA-NONOate, overnight cultures were diluted 1:50 into 200µl fresh LB, in the presence or absence of 2.5mM DETA-NONOate. Non-fluorescence WT IP2666 was grown alongside reporter strains for all experiments to determine the baseline fluorescence of Yptb grown under the indicated conditions.

### Fluorescence detection, plate reader

Bacterial samples were quantified directly in black-walled, clear bottom, 96 well plates. Optical density (A_600nm_) was used to approximate cell number by subtracting values from blank media control wells. For Yptb experiments, TIMER fluorescence was detected with 560nm excitation/610nm emission, and GFP fluorescence was detected with 480nm excitation/520nm emission, using a Synergy H1 microplate reader (Biotek). Fold change represents fluorescence in reporter strains relative to the average value for non-fluorescent WT Yptb. For *E. coli* experiments, TIMER signal was also detected with a spectral curve, with excitation at 480nm, and emission detected between 500nm-600nm.

### Fluorescence microscopy: bacteria

To visualize individual bacterial cells, 100µl of each sample was taken from 96 well plates and pelleted, resuspended in 100µl 4% PFA, and incubated at 4° C for 30 minutes for fixation. PFA was removed and bacteria were resuspended in 25µl PBS prior to imaging. Agarose pads were prepared to immobilize bacteria for imaging, by solidifying a thin layer of 25µl 1% agarose in PBS between a microscope slide and coverslip. Once solidified, coverslips were removed, bacteria (8µl) were added, coverslips were replaced, and bacteria were imaged with the 63x oil immersion objective, using a Zeiss Axio Observer.Z1 (Zeiss) fluorescent microscope with Colibri.2 LED light source and an ORCA-R^2^ digital CCD camera (Hamamatsu). Volocity image analysis software was used to specifically select individual bacterial cells and quantify the fluorescent signal associated with each cell.

### Murine model of systemic infection

Six to 8-week old female C57BL/6 mice were obtained from Jackson Laboratories (Bar Harbor, ME). All animal studies were approved by the Institutional Animal Care and Use Committee of Johns Hopkins University. Mice were injected intravenously into the tail vein with 10^3^ bacteria for all experiments (22, 26, 28). At the indicated timepoints post-inoculation (p.i.), spleens were removed and processed. Intact tissue was fixed for fluorescence microscopy. Tissue was homogenized to quantify CFUs.

### Fluorescence microscopy: host tissues

Spleens were harvested and immediately fixed in 4% PFA in PBS for either 3 hours at room temperature or 24 hours at 4° C. Tissues were frozen-embedded in Sub Xero freezing media (Mercedes Medical) and cut by cryostat microtome (Microm HM 505e) into 10µm sections. To visualize reporters, sections were thawed in PBS, stained with Hoechst at a 1:10,000 dilution, washed in PBS, and coverslips were mounted using ProLong Gold (Life Technologies). In Fig 3, neutrophils were also detected by immunofluorescence microscopy; sections were thawed, permeabilized with ice-cold 100% methanol and blocked in 2% BSA in PBS. Neutrophils were detected using a rat anti-mouse Ly6G unconjugated antibody (BD) and goat anti-rat Alexa 350-conjugated secondary antibody (Invitrogen). Sections were incubated with primary antibodies, washed, incubated with secondary antibodies, washed, and coverslips were mounted with ProLong Gold. Tissue was imaged as described above, with an Apotome.2 (Zeiss) for optical sectioning.

### qRT-PCR to detect bacterial transcripts in broth-grown cultures

Bacterial cells were grown in the presence or absence of DETA-NONOate for 4 hours, pelleted, resuspended in Buffer RLT (QIAGEN) + ß-mercaptoethanol, and RNA was isolated using the RNeasy kit (QIAGEN) according to manufacturer’s protocol. DNA contamination was eliminated using the DNA-free kit (Ambion). RNA was reverse transcribed using M-MLV reverse transcriptase (Invitrogen), in the presence of the RNase inhibitor, RnaseOUT (Invitrogen). Approximately 30 ng cDNA was used as a template in reactions with 0.5 µM of forward and reverse primers (22) and SYBR Green (Applied Biosystems). Control samples were prepared that lacked M-MLV, to confirm DNA was eliminated from samples. Reactions were carried out using the StepOnePlus Real-Time PCR system, and relative comparisons were obtained using the ΔΔC_T_ or 2^-ΔCt^ method (Applied Biosystems). Kits were used according to manufacturers’ protocols.

### Image analysis

Volocity image analysis software was used to quantify microcolony areas and fluorescence as previously described (22). Image J was used to quantify the signal intensity of each channel at the centroid and periphery of each microcolony, to generate relative signal intensities of fluorescent reporters, as previously described (22, 28). Thresholding was used to define the total area and centroid of each microcolony, and 0.01 pixel^2^ squares were selected to calculate values. Peripheral measurements depict bacteria in contact with host cells (22, 28).

### Statistical analyses for correlations

To determine if fluorescent signals were correlated, we performed linear regression analysis and are depicting best-fit lines. R^2^ values are shown to depict whether there is a linear correlation between the two fluorescent signals, however strength of correlation was assessed based on whether the slope of the best fit line was significantly non-zero. Low R^2^ values for many of the correlations are due to significant differences in the magnitude of fluorescent signals; TIMER_42_ signal is low relative to transcriptional fluorescent reporters. Additional statistical analyses were performed as described in figure legends.

## Author Contributions

Conceptualization: PP, BJO, EA, KMD; Formal Analysis: PP, KMD; Funding Acquisition and Supervision: KMD; Investigation: PP, BJO, KMD; Methodology: PP, BJO, EA, KMD; Writing – Original Draft Preparation: PP, KMD; Writing – Review & Editing: PP, BJO, EA, KMD.

## Acknowledgments

We thank the members of the Davis lab, who provided feedback and suggestions during the final steps of manuscript preparation. We also thank Dr. Ralph R. Isberg for guidance and feedback throughout this project. The authors of this manuscript declare no conflicts of interest. This work was supported by a NIAID K22 Career Transition Award (1K22AI123465-01).

---

**Supplemental Fig 1:**
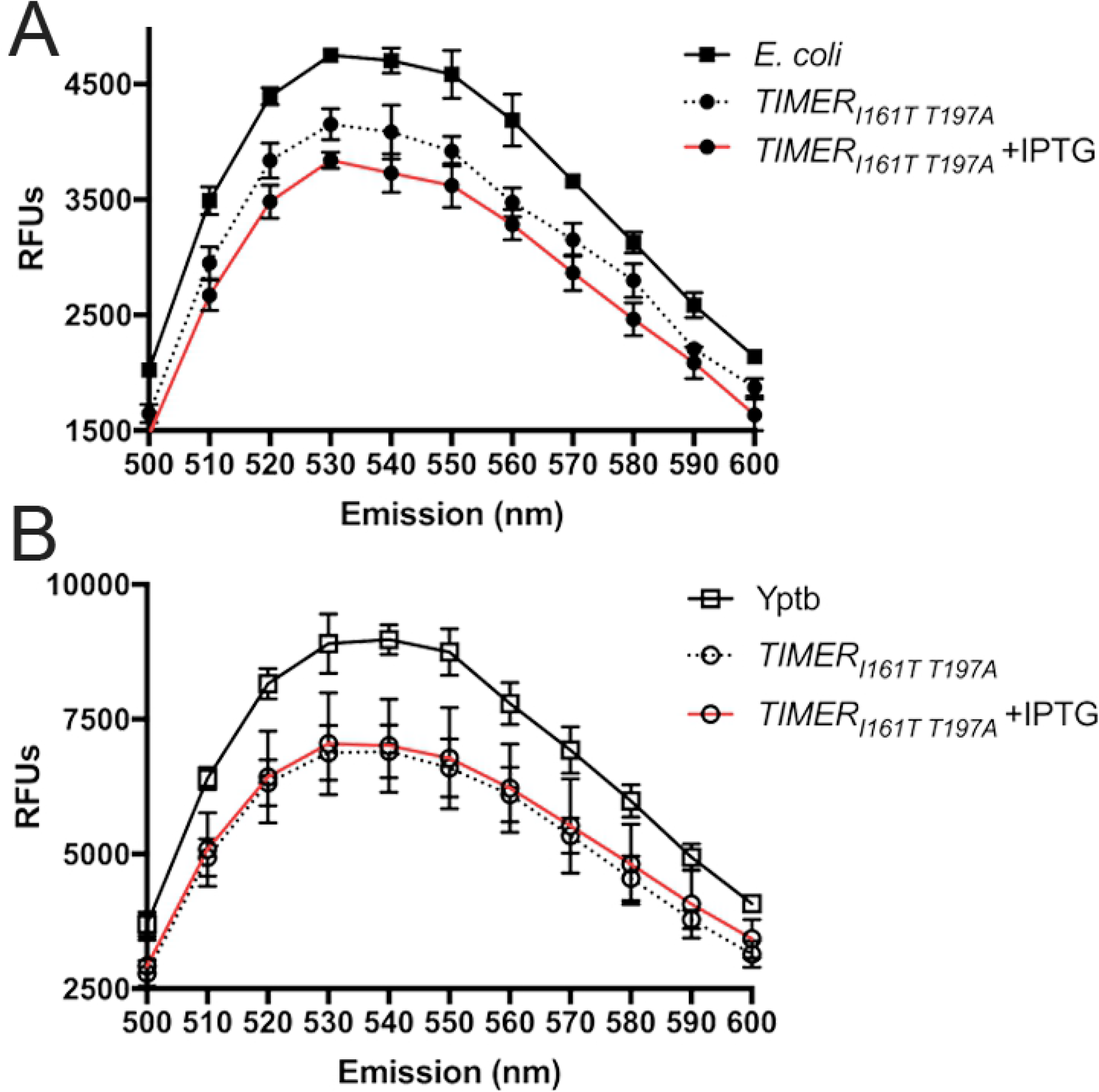
*TIMER*_*I161T T197A*_ lacks fluorescence. The indicated strains contain the *TIMER*_*I161T T197A*_ variant in the low copy vector, pMMB67EH. Strains were grown at 37° C in 96 well plates with shaking for 24h. Fluorescence was detected with a spectral curve with excitation at 480nm and emission detected between 500nm-600nm. Relative fluorescence units (RFUs) represent raw values with media only background subtracted, data represents mean of three biological replicates. A) *E. coli* strains: *E. coli* is the non-fluorescent DH5αλpir *E. coli* parent strain. B) *Y. pseudotuberculosis* strains: Yptb is the non-fluorescent WT IP2666 parent strain.

**Supplemental Fig 2:**
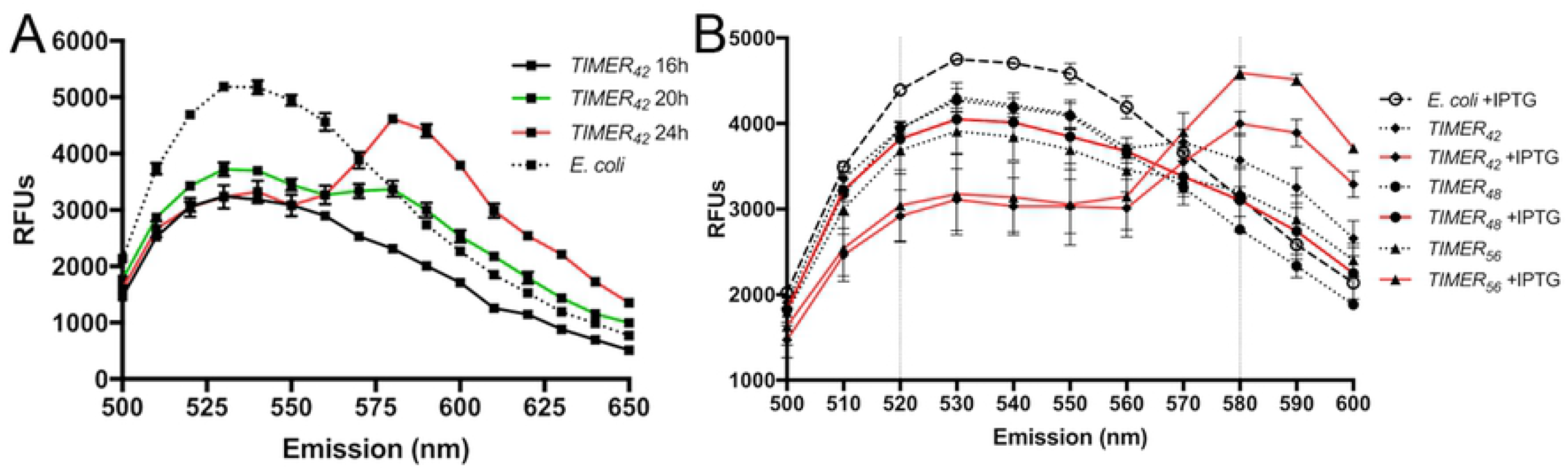
Red fluorescence emerges in TIMER variants. *E. coli* strains (XLI-BLUE background) contain the indicated TIMER variants, expressed from the high copy pUC18 plasmid. Strains were grown for the indicated times at 37° C in 96 well plates with shaking. Fluorescence was detected with a spectral curve with excitation at 480nm and emission detected between 500nm-600nm. Relative fluorescence units (RFUs) represent raw values with media only background subtracted, data represents mean of three biological replicates. A) TIMER_42_ variants were grown alongside the non-fluorescent WT *E. coli* strain in the presence of IPTG for the indicated times. *E. coli*: 24h growth. B) TIMER variants grown alongside the non-fluorescent WT *E. coli* strain, in the presence or absence of IPTG for 24h. Vertical dotted lines: 520nm (green fluorescence) and 580nm (red fluorescence).

**Supplemental Fig 3:**
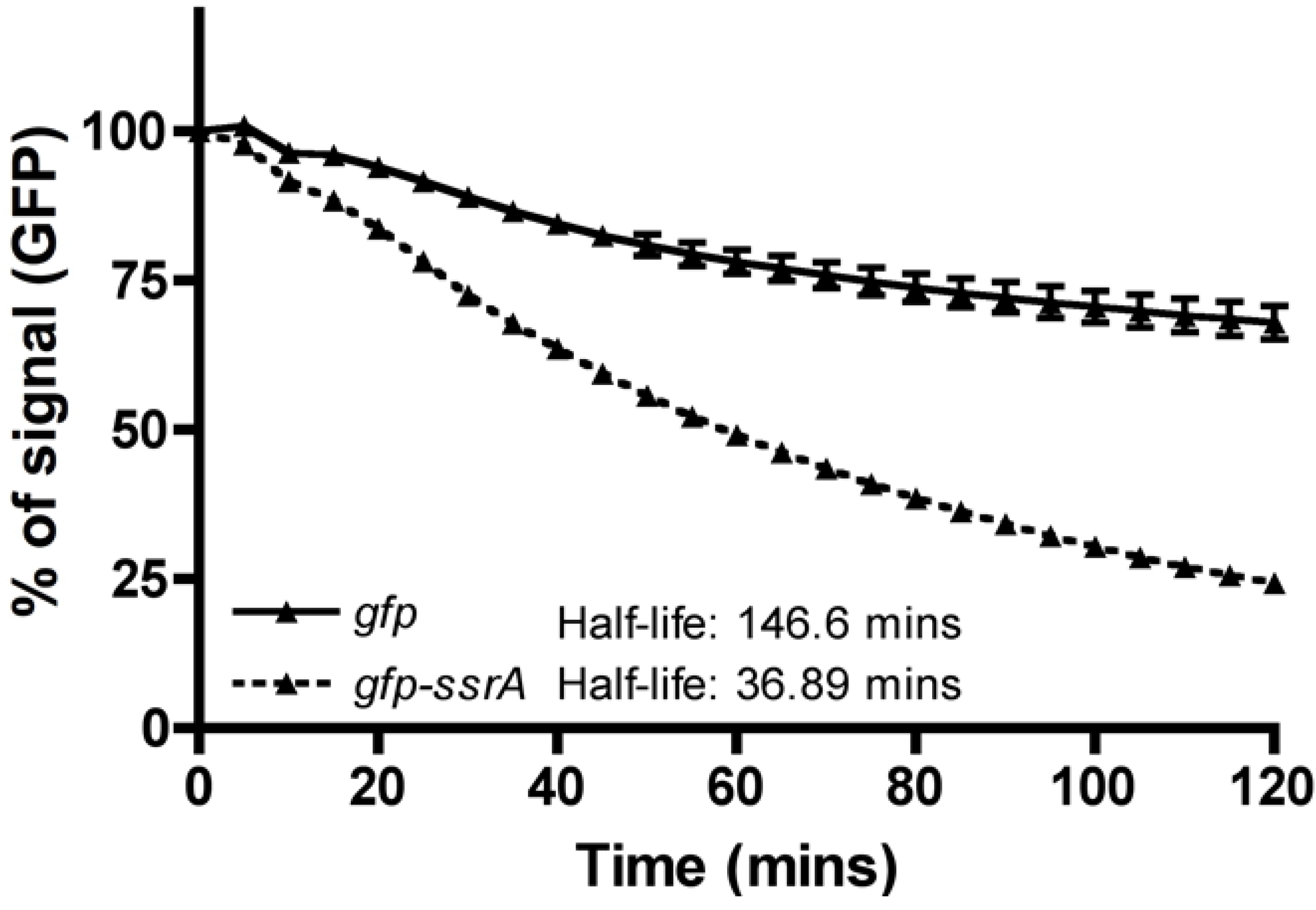
Stability of GFP and GFP-ssrA. WT IP2666 containing IPTG-inducible *gfp* or *gfp-ssrA* were grown overnight (16h) in the presence of IPTG. Cells were pelleted and resuspended in PBS, and kanamycin was added to inhibit additional protein translation. GFP signal was detected with 480nm excitation/520nm emission every 5 minutes for 120 minutes in a plate reader, to detect the half-life of each fluorescent protein by the one-phase exponential decay equation. Fluorescent values are expressed as % of signal (GFP), where time 0 equals 100% signal intensity. Data represents mean of three biological replicates.

